# The role of catalytic and regulatory domains of human PrimPol in DNA binding and synthesis

**DOI:** 10.1101/2023.01.09.523353

**Authors:** Elizaveta O. Boldinova, Andrey G. Baranovskiy, Diana I. Gagarinskaya, Alena V. Makarova, Tahir H. Tahirov

## Abstract

Human PrimPol possesses DNA primase and DNA polymerase activities and restarts stalled replication forks protecting cells against DNA damage in nuclei and mitochondria. The zinc-binding motif (ZnFn) of the C-terminal domain (CTD) of PrimPol is required for DNA primase activity but the mechanism is not clear. In this work, we biochemically demonstrate that PrimPol initiates *de novo* DNA synthesis in *cis-*orientation, when the N-terminal catalytic domain (NTD) and the CTD of one molecule take part in catalysis. The modeling studies revealed that PrimPol uses a similar mode of initiating NTP coordination as the human primase. The ZnFn motif residue Arg417 is required for binding the 5’-triphosphate group that stabilizes the PrimPol complex with a DNA template-primer. We found that PrimPol is able to efficiently initiate DNA synthesis in the absence of the link between the two domains. The ability of the NTD alone to prime DNA synthesis and a regulatory role of the RPA-binding motif in the modulation of PrimPol binding to DNA are also demonstrated.

## INTRODUCTION

Human primase polymerase PrimPol, belonging to the archaeo-eukaryotic superfamily, was described in 2013 (1–3). PrimPol is encoded by the *PRIMPOL* gene located on chromosome 4 at locus 4q35.1, and is a protein consisting of 560 amino acid residues with a molecular mass of 65 kDa (1, 4). In contrast to the human replicative primase heterodimer PriS/PriL that synthesizes 9-mer RNA primers (5), PrimPol engages in *de novo* synthesis using deoxyribonucleotides, resulting in DNA primers that do not require removal.

PrimPol is present in both the nucleus and mitochondria (1). PrimPol primase activity allows for replication re-initiation at DNA sites containing damage or at secondary structures that block high-fidelity DNA polymerases (6–9). Lack of PrimPol in the cell leads to a slowing down of replication, chromosomal aberrations, and increased sensitivity to various DNA-damaging agents (1–3, 10, 11).

PrimPol is composed of two domains: an N-terminal AEP-like catalytic domain (NTD), and a C-terminal domain (CTD) similar to the UL52 primase of the herpes simplex virus (Figure 1) (4, 12). Two crystal structures of the human NTD (residues 1 to 354) in complex with a primer/undamaged DNA template and with a primer/DNA template with 8-oxo-G and incoming nucleotides have been deciphered (PDB ID: 5L2X; PDB ID: 7JK1) (13, 14). The active site of the NTD is formed by the N-helix (residues 1-17) and two modules: ModN (residues 35-105) and ModC (residues 108-200 and 261-348). Conservative motifs I (DxE) and III (hDh) of ModC contain the key catalytic amino acid residues Asp114/Glu116 and Asp280 involved in the coordination of Me^2+^ ions, while motif II (SxH) contains Ser167 and His169 residues that bind the incoming nucleotide (13). Mutations of Asp114/Glu116, Asp280, and His169 residues result in loss of DNA polymerase and primase activities of human PrimPol (1–3, 15).

**Figure 1.**
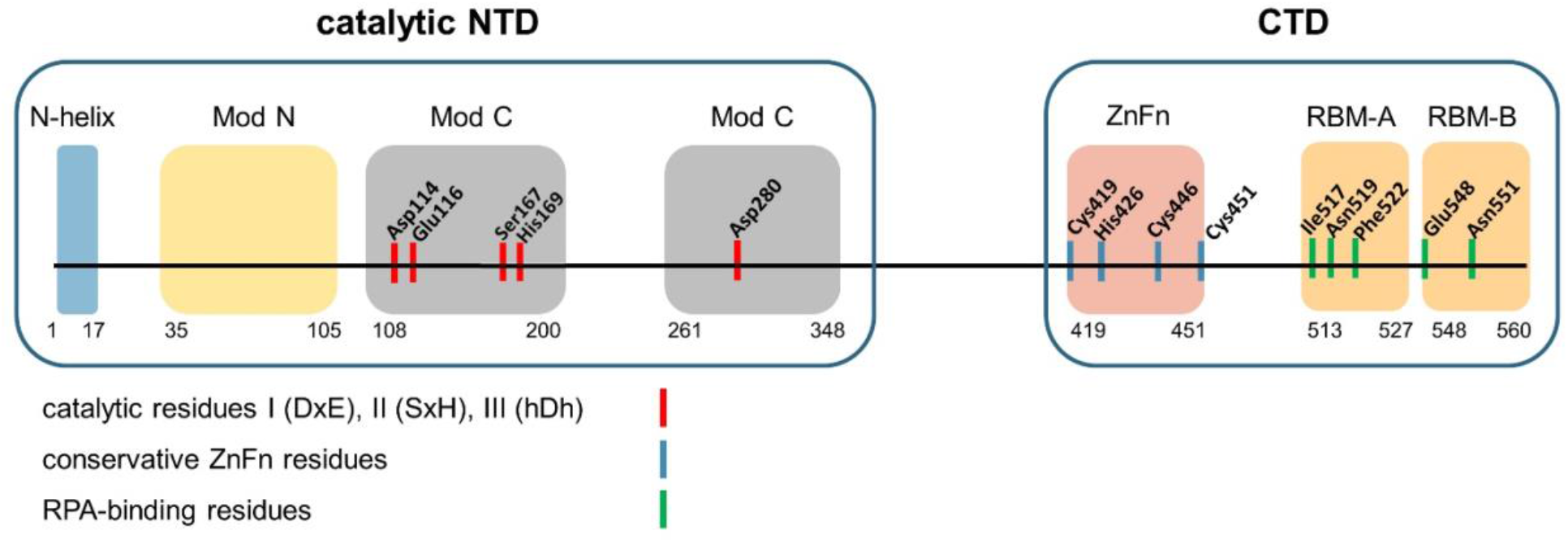
Domain organization of PrimPol. The positions of the catalytic residues coordinating Me^2+^ ions (Asp114, Glu116, Asp280), the conservative C-H-C-C ZnFn motif, and the RPA-binding motif residues are indicated.

Conserved motifs Ia (RQ) (residues Arg47 and Gln48) and Ib (QRhY/F) (residues Gln75, Arg76, and Tyr78) of the ModN module form contacts with a DNA template (13). Mutations of the Arg47 and Arg76 residues significantly reduce the DNA polymerase and primase activities of PrimPol (16). The N-helix is connected to ModN by a long flexible linker (residues 18-34) and interacts with the DNA template strand, by making a few contacts with the major groove (13). Unlike DNA-polymerases, the NTD of PrimPol is almost completely devoid of contacts with a primer, and its active site has room to accommodate the initiating NTP to form a dinucleotide during *de novo* DNA synthesis (13).

The CTD domain contains a conserved C-H-C-C zinc finger (ZnF) motif that coordinates the Zn^2+^ ion (∼372–487 residues harboring Cys419, His426, Cys446, Cys451) (7, 12, 17). It was shown that ZnFn is required for *de novo* DNA synthesis but not for DNA polymerase activity. Deletion of ZnFn and mutations of the conservative residues Cys419 and His426 that coordinate zinc disrupt PrimPol primase activity while retaining DNA polymerase activity (3, 7, 12, 17).

Amino acid residues 201 to 260 of the ModC module represent an unstructured region and presumably play a regulatory role. Residues 226 to 232 form contacts with the accessory protein PolDIP2 (18, 19). Residues 60-70 of the ModN domain of PrimPol form the second binding site for the replicative factor PolDIP2 (19). The CTD contains the RPA-binding domain (RBD) harboring two negatively charged RPA-binding motifs (RBM): RBM-A (residues 513 – 527), and RBM-B (residues 546 – 560) (20).

The detailed mechanism of PrimPol primase activity is not fully understood. Structures of the full-length PrimPol and the initiation complex of PrimPol with a DNA template and a dinucleotide have not been deciphered. However, some important information about the mechanism of PrimPol primase activity has been obtained from biochemical studies. PrimPol prefers to initiate DNA synthesis on the 3’-GTC-5’ sequence with cryptic G (1, 21), and its activity is dependent on cofactor metal ions. PrimPol exhibits DNA polymerase activity in the presence of Mn^2+^ and Mg^2+^ ions, but primase activity is stimulated exclusively by Mn^2+^ ions (15, 22). It has been suggested that after binding and recognizing the preferred DNA site, PrimPol binds the first nucleotide at the elongation site, which is stabilized by Mn^2+^ ions (17). It is assumed that binding of the second nucleotide (preferably ATP) occurs at the initiation site, followed by catalysis and formation of the rA-dG dinucleotide.

According to the suggested model, there is a division of work between the catalytic subunit/domain and the accessory subunit/domain (often containing ZnF or an iron-sulfur cluster) in primases (5). In human primase, the small catalytic PriS subunit is responsible for catalysis, whereas the flexibly tethered large accessory PriL subunit is responsible for the binding of a template and initiating NTP. The interaction of PriL with the RNA/DNA hybrid involves the CTD and the 5′-triphosphate of an RNA primer (5). In a similar way, dinucleotide formation with adenosine triphosphate is 16 times more efficient than with adenosine di- and monophosphates, suggesting that a triphosphate group at the 5ʹ-initiator nucleotide is required for *de novo* DNA synthesis by PrimPol (17). Moreover, the elongation of a dinucleotide by PrimPol is possible only in the presence of a 5ʹ-triphosphate on the initiating nucleotide (17). Deletion of the ZnFn suppresses dinucleotide formation with ATP and impedes further primer elongation, which suggests the ZnFn motif plays a role in the stabilization of ATP at the 5ʹ-initiator site and/or in the coordination of cryptic Gua of a DNA template (17, 23).

Due to bi-modal organization, primases can operate in the *cis-* or *trans-*orientation. The *cis-*mechanism implies that the catalytic subunit/domain and accessory subunit/domain of the same molecule are involved in catalysis. In the *trans-*orientation, the accessory unit of one primase molecule binds DNA and/or the initiator nucleotide, while dinucleotide formation takes place in the active site of another molecule. DNA primase of phage T7 (24) and primase RepB (25) are able to initiate primer synthesis in the *trans*-mode, while human primase is not (26, 27). The mechanism of PrimPol operation is currently unknown.

In the present work, we analyzed the mechanism of PrimPol primase activity using its variants that selectively disrupt catalytic DNA-synthetic function or only priming activity, as well as the individual catalytic domains of PrimPol. We show that PrimPol performs primer synthesis in the *cis-*orientation, when the NTD catalytic and CTD regulatory domains of the same molecule take part in catalysis. We also demonstrate a regulatory role of the CTD in template-primer binding by PrimPol.

## MATERIAL AND METHODS

### Protein purification

The wild-type PrimPol and all mutant variants fused at the N-terminus with a SUMO-HIS_10_ tag were purified from *E. coli* cells as described (28).

### DNA substrates

Oligonucleotides were synthesized by Eurogene and Syntol (Moscow, Russia). To prepare the DNA substrate for testing DNA polymerase activity, Primer-18 was 5′-labeled with [γ-^32^P]-ATP by T4 polynucleotide kinase (SibEnzyme, Russia) and annealed to the corresponding unlabeled Template-55 at a molar ratio of 1:1.1, heated to 75 ºC, and slowly cooled down to 24 ºC. The sequences of the oligonucleotides used in this study are shown in Table 1.

**Table 1.**
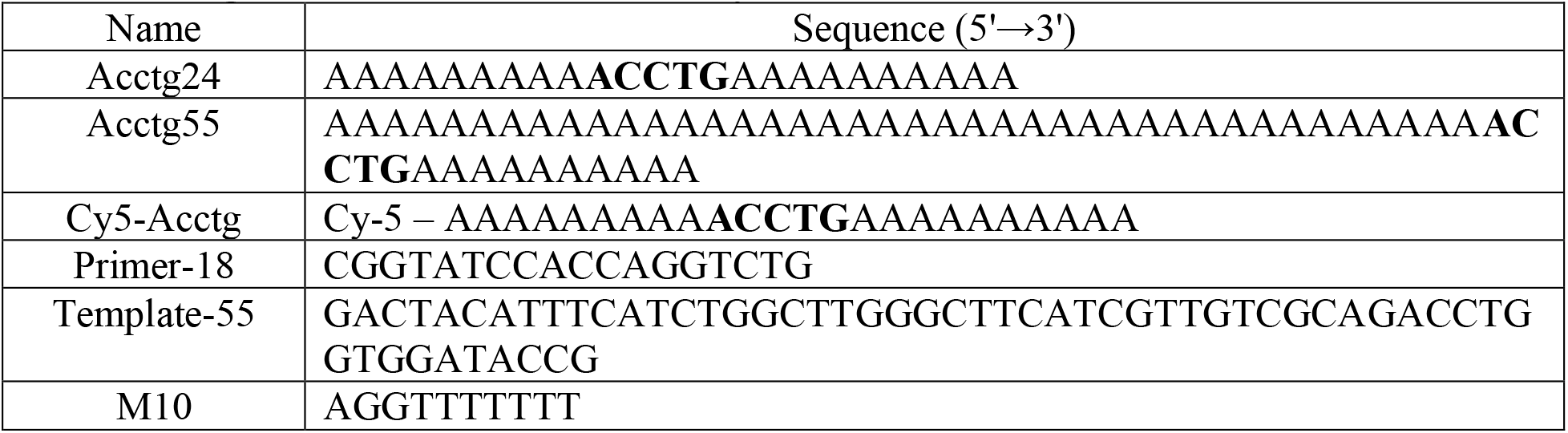
Oligonucleotides used in the study.

Oligonucleotide “p-p-p-13” containing the 5ʹ-terminal adenosine triphosphate was synthesized enzymatically *de novo* using the wild-type PrimPol. 600 μL of reaction mixture (aliquoted in 6 tubes at 100 μl) contained 40 mM HEPES pH 7.0, 8% glycerol, 50 μg/mL BSA, 1 mM MnCl_2_, 10 mM MgCl_2_, 10 μM “Acctg24” template oligonucleotide, 200 μM dGTP and dTTP, 10 μM dATP, 1 mM rATP, and 5 μM PrimPol. The reactions were incubated for 4 h at 30 ºC, and 2.5 μM fresh PrimPol was additionally added to the reaction every hour. DNA was precipitated from solution by ethanol. Three volumes of ice-cold 96% ethanol, 300 mM NaCl, 10 mM MgCl_2_, and 1 μg/μl glycogen were added to 1 volume of solution and incubated overnight at -20 ºC. After centrifugation for 30 min at 14 000 g and 0 ºC, the precipitate was washed with 70% ice-cold ethanol, dried, and dissolved in 35 μl of H_2_O. Next, an equal volume of loading formamide mixture (95% formamide, 20 mM EDTA, 0.05% bromophenol blue) was added and tubes were heated at 95 ºC for 5 min. The synthesized product and template were separated on denaturing 16% PAGE with 7 M urea in 1xTBE. DNA imaging was carried out by the UV shadowing of DNA spots on a fluorescent thin-layer chromatographic Silufol plate (Figure S1). Bands corresponding to 12-13 nt oligonucleotides were excised from the gel, crushed, and extracted in 1 ml of extraction buffer (500 mM NH_4_Ac, 10 mM MgAc_2_, 1 mM EDTA, pH 7.0) overnight with shaking. DNA was precipitated from solution by ethanol and dissolved in 35 μl of H_2_O.

### Primase reactions

DNA primase activity was tested in 6 μl reaction mixtures containing 40 mM HEPES pH 7.0 (or 7.4), 8% glycerol, 50 μg/μl BSA, 1 mM MnCl_2_, 2 μM unlabeled oligonucleotide DNA substrate, 200 μM each of dGTP, dCTP, and dTTP or dGTP alone, 10 μM dATP, 30 nM [^γ-32^P]-ATP, and 2 μM PrimPol. Test tubes were preincubated on ice for 5 min and reactions were started by dNTP and incubated at 30 ºC for 60 min or for the indicated time intervals. The reactions were stopped by adding an equal volume of loading formamide mixture. Experiments were repeated twice for each protein preparation.

### DNA polymerase reactions

Primer extension reactions were performed in 20 μl reaction mixtures containing 20 nM radioactively labeled oligonucleotide substrate, 200 μM of each dNTP, 40 mM HEPES pH 7.0, 8% glycerol, 50 μg/ml BSA, 100 nM PrimPol, and 10 mM MgCl_2_ or 1 mM MnCl_2_. Test tubes were preincubated on ice for 5 min and reactions were started by dNTP and incubated at 37 ºC for the indicated times. The reactions were stopped by adding an equal volume of loading formamide mixture. Experiments were repeated twice for each protein preparation.

### EMSA

Binding of PrimPol to the ^32^P-labeled Template55/Primer-18 substrate, the fluorescently labeled ssDNA substrate Cy5-Acctg24, and the DNA template-primer substrate with p-p-p-13 annealed to Cy-Acctg24 was performed in a 20 μl reaction mixture containing 40 mM HEPES pH 7.0, 30 mM KCl, 1 mM DTT, 8% glycerol, 0.1 mg/ml BSA, 1 mM MnCl_2_, 300 nM DNA substrate, and 250-1200 nM PrimPol. Reactions were supplemented with 1 mM ATP and 200 μM of each dNTP as indicated in the figure legends. The reactions were incubated at 24 ºC for 20 min and placed on ice. Next, 5 µl of 50% glycerol with 0.05% bromophenol blue was added, and the PrimPol:DNA complex was separated from free DNA in a 5% native PAAG in 0.5x Tris-glycine buffer (12.5 mM Tris, 96 mM glycine, pH 8.3) at 10 V/cm and at 4 ºC. The gel was visualized on a Typhoon 9400 (GE Healthcare, USA).

## RESULTS

### PrimPol operates in *cis*-orientation

To analyze the mechanism of primase activity, we replaced the key amino acid residues of the NTD and CTD of the full-length PrimPol: D114A substitution of the Asp114 residue coordinating Me^+^ ions (PrimPol_D114A_), and R417A and R424A substitutions of conserved ZnFn motif residues that presumably bind the 5’-triphosphate of initiating ATP (PrimPol_R417A_ and PrimPolR_424A_) (Figure 2A). In addition to these separation-of-function mutations, we also analyzed variants of PrimPol with deletions of the NTD (PrimPol_363-560_) or CTD (PrimPol_1-363_ and PrimPol_1-363CD_) and RBM (PrimPol_1-475_) (Figure 2A).

**Figure 2.**
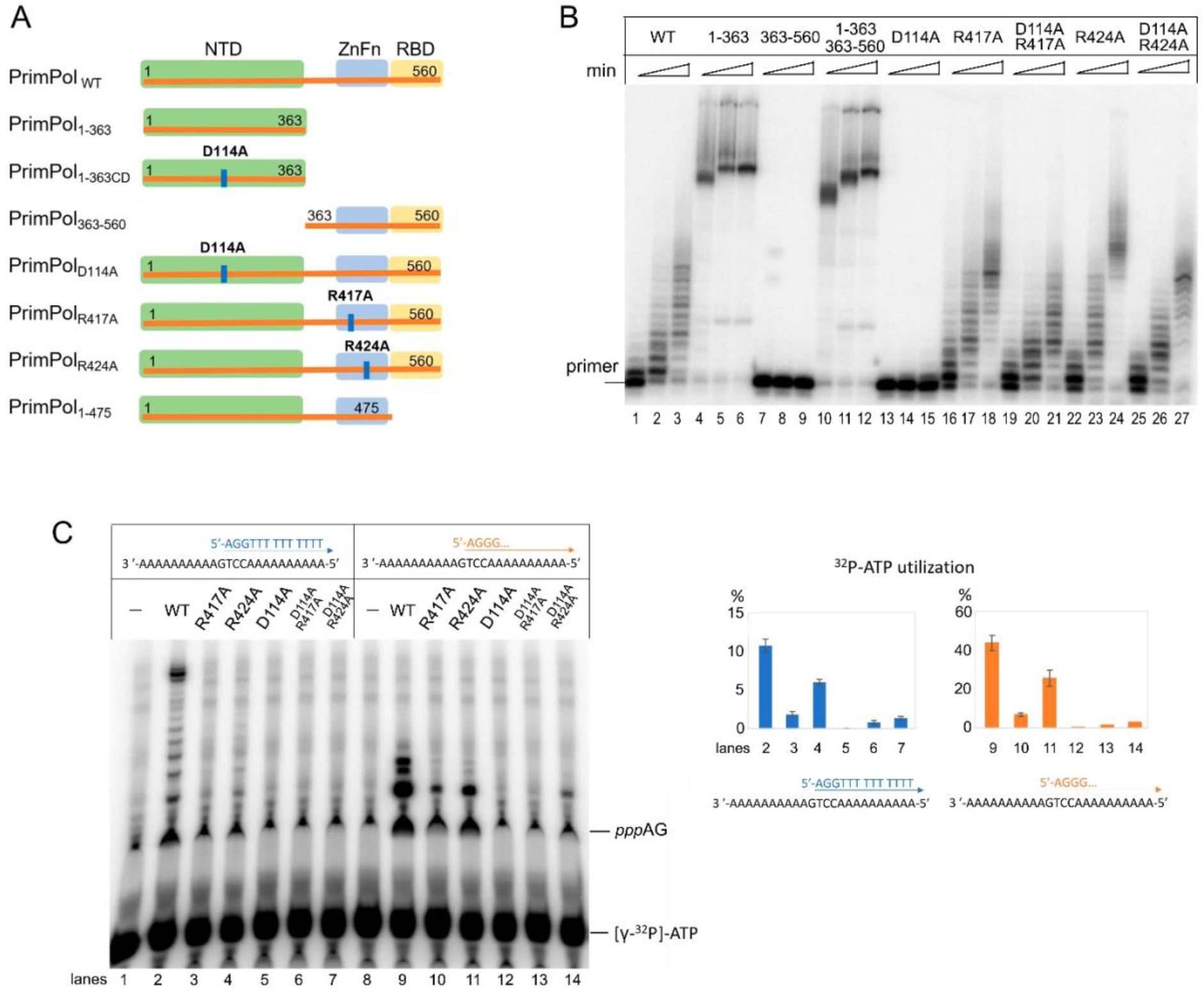
The DNA polymerase and primase activity of PrimPol variants with separation-of-function point mutations. **A**. The scheme of PrimPol point mutations and deletions. **B**. The DNA polymerase activity of PrimPol mutant variants. Reactions were incubated with the ^32^P-labeled Template55/Primer-18 substrate for 2–30 min; pH 7.0 and 10 mM Mg^2+^. **C**. The DNA primase activity of PrimPol variant. Reactions were incubated in the presence of [γ-^32^P]-ATP, ATP, dGTP, and dTTP (lanes 1–7) or [γ-^32^P]-ATP, ATP, and dGTP (lanes 8–14); pH 7.0 and 1 mM Mn^2+^.

The DNA polymerase activity (Figure 2B), total DNA primase activity and dinucleotide formation (Figure 2C) of PrimPol variants and their combinations were analyzed. The D114A substitution resulted in the loss of any catalytic activity (Figure 2B, lanes 13–15 and Figure 2C, lanes 5,12). The full-length PrimPol variants with ZnFn substitutions R417A and R424A retained DNA polymerase activity while their primase activity was severely affected (Figure 2C, lanes 3,4,10,11). These data indicate that R417 and R424 are important only upon priming of DNA synthesis. This is consistent with previous studies showing the key role of the CTD in DNA synthesis initiation but not elongation (3, 7, 12, 17, 23).

In the case of the *trans*-mechanism of PrimPol, mixing in one reaction the two mutant forms PrimPol_D114A_ and PrimPol_R417A_ or PrimPol_D114A_ and PrimPolR_424A_ would result in the formation of functional dimer molecules carrying one functional catalytic NTD domain and one functional regulatory CTD. As a result of the cooperation between the functional domains of different mutant forms, the restoration of DNA-primase activity is be expected. However, when mutant variants PrimPol_D114A_ and PrimPol_R417A_ or PrimPol_D114A_ and PrimPol_R424A_ were mixed in the reaction, no restoration of the DNA-primase activity occurred. On the contrary, a slight decrease of all activities was observed in these reactions compared to reactions with PrimPol_D117A_ and PrimPol_R424A_ variants (Figure 2). The inhibition may result from the competition of catalytically active ZnFn-mutant variants with the catalytically inactive PrimPol_D114A_ variant for a DNA substrate. These results are consistent with the model of the *cis*-orientation when the catalytic NTD and regulatory CTD of the same PrimPol molecule are involved in DNA synthesis.

### The NTD retains weak primase activity and cooperates with the separated CTD

Remarkably, the full-length PrimPol_R417A_ and PrimPol_R424A_ variants demonstrated levels of DNA polymerase activity similar to the wild-type enzyme (Figure 2B, lanes 16–18, 22–24), but deletion of the CTD (variant PrimPol_1-363_) dramatically increased the DNA polymerase activity (Figure 2B, lanes 4–6). Moreover, separate NTD retained weak DNA primase activity (Figure 3A).

**Figure 3.**
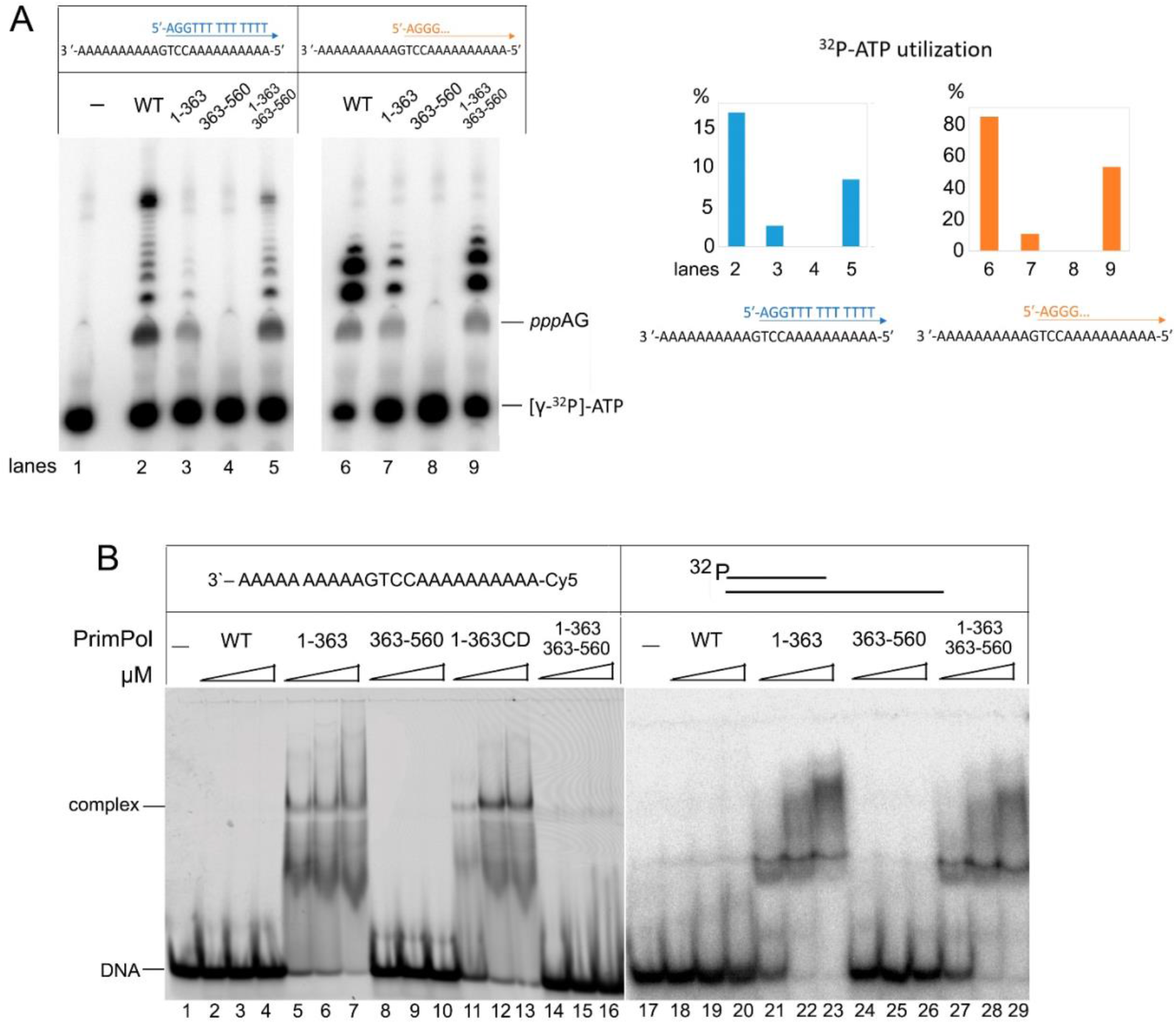
The DNA polymerase and primase activity of PrimPol variants with separate NTD and CTD. **A**. The DNA primase activity of PrimPol variants. Reactions were incubated in the presence of [γ-^32^P]-ATP, ATP, dGTP, and dTTP (lanes 2–5) or [γ-^32^P]-ATP, ATP, and dGTP (lanes 6–9); pH 7.0 and 1 mM Mn^2+^. **B**. EMSA of ssDNA Cy5-Acctg and the ^32^P-labeled Template55/Primer-18 binding by PrimPol variants. Reactions were incubated with 1, 2, or 3 µM PrimPol in the presence of 500 nM DNA and 1mM Mn^2+^, pH 7.0.

Interestingly, the DNA primase activity of the PrimPol_1-363_ variant was higher at pH 7.5 compared to 7.0 (Figure 2S). This is contrary to the full-length PrimPol showing higher DNA polymerase (29) and DNA primase (Figure 2S) activities at pH 7.0 versus pH 7.5. The difference may be due to different isoelectric points of PrimPol (5.14) and NTD (7.61) or to a different mode of initiating NTP coordination in the absence of CTD. The absence of impurities and an extrinsic primase activity in the NTD preparation was verified using the catalytically inactive PrimPol_1-363CD_ variant harboring the D114A substitution (Figure S3) and mass-spectrometry.

At pH 7.0, PrimPol_1-363_ shows weak primase activity, while PrimPol_363-560_ has no catalytic activity (Figure 3A). When NTD and CTD were mixed together, the primase activity was partially restored (Figure 3A, lanes 5 and9). This result points to cooperation of the two separated domains during initiation of DNA synthesis and to the possible interaction between them. Indeed, with a broken linkage between NTD and CTD, only a relatively stable interaction of these two domains could result in almost complete restoration of primase activity. The notion of interaction between the two PrimPol domains upon dinucleotide formation is supported by EMSA with single-stranded DNA (Figure 3B, lanes 14–16). Importantly, PrimPol_1-363_ makes a stable complex with DNA, while the full-length PrimPol and PrimPol_363-560_ do not bind ssDNA. Mixing NTD and CTD together resulted in the loss of PrimPol_1-363_ ability to bind ssDNA. These data suggests that the CTD of PrimPol might prevent efficient binding of the protein to DNA. The high affinity of PrimPol_1-363_ to DNA in the absence of CTD may explain its high DNA polymerase activity (Figure 2B). Thus, CTD demonstrates negative regulation of ssDNA binding by NTD. This is the opposite of human primase, where PriS has very low affinity to DNA and CTD plays the main role in template-primer binding (5). However, CTD did not abolish the NTD binding to the template-55/primer-18 (Figure 3B, lanes 27–29). The absence of a negative effect of CTD on the NTD/dsDNA complex can be explained by the higher PrimPol affinity to a primed DNA template.

### The ZnFn Arg417 and Arg424 are required for the interaction with the 5’-triphosphate

During *de novo* DNA synthesis, PrimPol predominantly uses ATP (or dATP) as the initiator nucleotide, and the ZnFn motif of the CTD plays a key role in binding and incorporation of ATP by an unknown mechanism (17, 23). We propose that the conservative ZnFn residues Arg417 and/or Arg424 may play a role in binding the 5’-triphosphate of the initiating ATP, which becomes the first nucleotide of the primer. We selected these arginines because they are the most conservative in the ZnFn motif (7). By analogy, in PriL of human primase, Arg302 and Arg306 interact with the 5’-triphosphate of a primer and play the key role in primase activity (30). Indeed, the R417A and R424A substitutions disrupted the DNA primase activity of PrimPol in this work. To study the possible role of these arginines in the 5’-triphospate binding, a DNA substrate with a primer containing adenosine triphosphate at the 5’-end was synthesized enzymatically using PrimPol (Figure 1S).

We have shown that the wild-type and mutant forms of PrimPol cannot efficiently bind the DNA duplex without a triphosphate at the 5’-end (Figure 4B, lanes 2–7). In contrast, PrimPol binds the DNA duplex with the 5’-triphosphate very efficiently (Figure 4B, lanes 8–9). Moreover, the R424A and R417A substitutions dramatically decreased the ability of PrimPol to bind DNA with the 5’-triphosphate (Figure 4B, lanes 10–13). These data indicate that PrimPol holds the primer 5’-end during all steps of DNA primer synthesis like a human primase.

**Figure 4.**
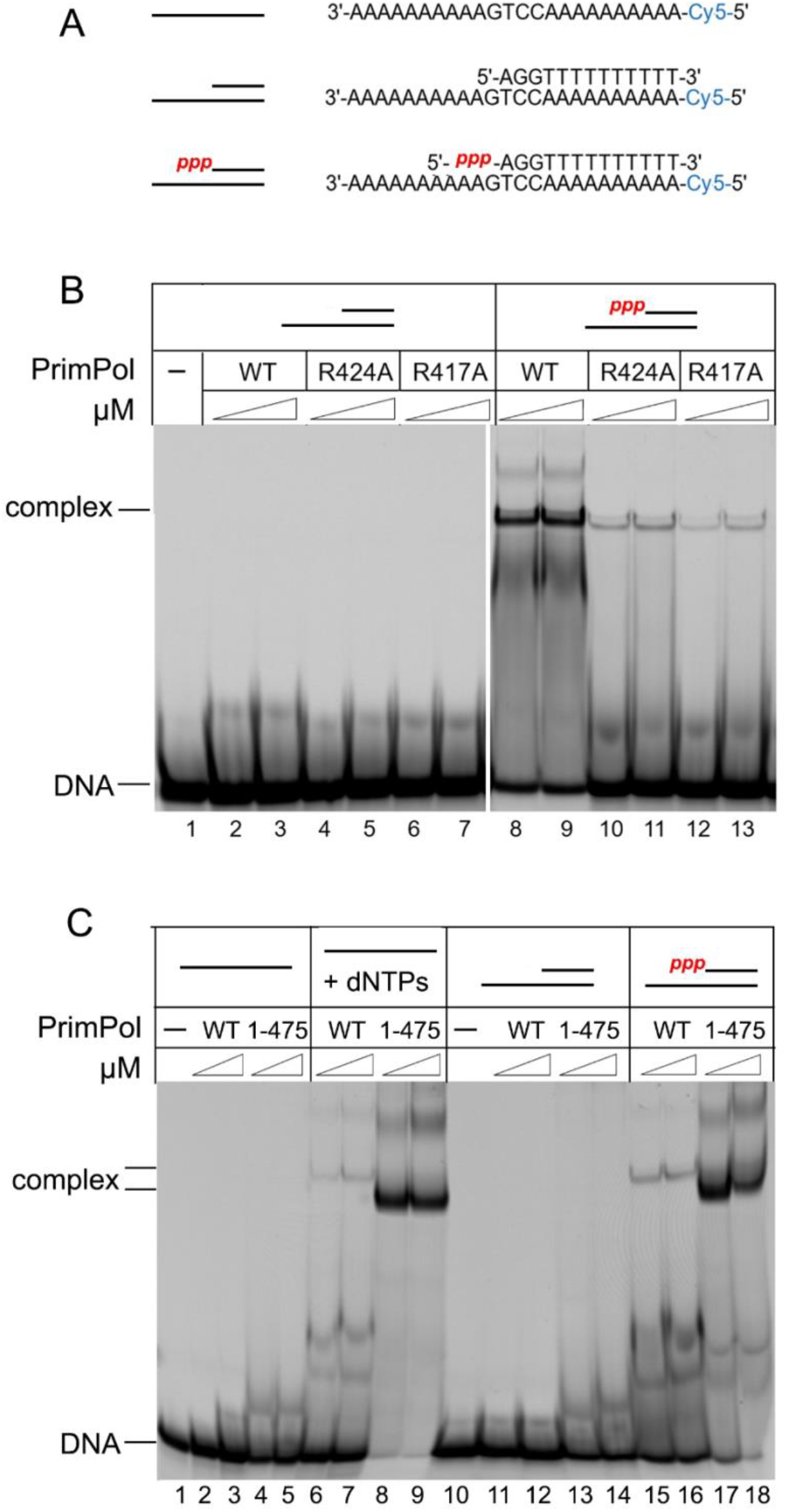
The role of 5’-triphospate and RBM in DNA binding of PrimPol. **A**. Structures of DNA substrates used in the study. **B**. EMSA of binding to DNA with 5’-triphospate by PrimPol ZnFn mutant variants. Reactions were incubated with 1 or 2 µM PrimPol in the presence of 300 nM DNA and 1mM Mn^2+^, pH 7.0. **C**. EMSA of binding to DNA by the PrimPol RBM mutant variant. Reactions were incubated with 1 or 2 µM PrimPol in the presence of 300 nM DNA and 1mM Mn^2+^, pH 7.0. ATP, dGTP, and dTTP were added to some reactions (lanes 6–9).

### The RBM modulates PrimPol binding to DNA

In addition to the ZnFn motif, the CTD contains another important element, the RPA-binding motif (RBM), which is rich in negatively charged a.a. (31). It can be assumed that this motif may affect the affinity of PrimPol to DNA by interacting with RPA. Indeed, the PrimPol_1-475_ variant with deletion of the RBM showed increased DNA binding efficiency to DNA with the 5’-triphosphate compared to the full-length protein (Figure 4C, lanes 17, 18). These data demonstrate the key role of the CTD and particularly the RBM motif in regulation of PrimPol binding to DNA. The mechanism of regulation of PrimPol activity may be due to the interaction of RPA or other proteins with the negatively charged RBM and neutralization of the excess charge, which promotes DNA binding by the enzyme and its activity.

As expected, DNA binding of the wild-type PrimPol and the PrimPol_1-475_ variant was more efficient on DNA with the 5’-triphosphate, which was synthesized and purified before analysis (Figure 4B, lanes 17, 18) or incorporated by PrimPol during *de novo* DNA synthesis directly in EMSA reactions (Figure 4B, lanes 6–9).

## DISCUSSION

### Human PrimPol initiates DNA synthesis in *cis-*orientation

In this work, we demonstrated that the full-length PrimPol operates in *cis*-orientation when the catalytic NTD and regulatory CTD of the same molecule are involved in *de novo* DNA synthesis. This mechanism is similar to that previously described for replicative human primase PriS/PriL (5). On the other hand, experiments with separated NTD and CTD revealed their ability to work in *trans-*orientation. We suggest that the actual test for the *trans-*mechanism of DNA synthesis priming is experiment involving point mutations (when a molecule with mutation in the CTD can complement primase activity of the NTD mutant). Experiments with CTD and NTD mixed together revealed their ability to cooperate without the covalent link between them, which points to non-covalent complex formation. The interaction between the catalytic and regulatory domains would stabilize the initiation complex composed of several weakly bound components: PrimPol, DNA template, and two NTPs.

### The role of the ZnFn motif in primase activity

The C-terminal ZnFn motif plays a key role in the primase activity of PrimPol (17, 23). In this work, we demonstrated that the triphosphate at the 5’ end of the primer is required for efficient binding of PrimPol to the template-primer. Our results support the findings of Martínez-Jiménez et al. (17) showing that PrimPol uses nucleotide triphosphate as the initiating nucleotide (di- and monophosphates significantly reduce primase activity).

Despite the key role of the ZnFn in primase activity, the NTD of PrimPol lacking the ZnFn retains weak primase activity. These data are in agreement with recent studies showing weak primase activity of the NTD of PrimPol (deletion 410–560 a.a.) (17) and CRISPR-associated primase-polymerases (32) in reaction with Mn^2+^. In an earlier study, the mutant N-terminal variant PrimPol_1-354_ completely lacked primase activity (12). Contrasting results may be related to sensitivity of the NTD activity to the low pH (7.0) widely used in assays and to the absence of Mn^2+^ ions in primase reaction. In addition, one NTD molecule may assist another by binding the initiating nucleotide during dinucleotide synthesis. Higher primase activity of NTD at pH 7.5 can be explained by more stable/optimal interaction between the two NTD molecules during dinucleotide formation.

We built the model of the ZnFn/template-primer complex using the coordinates of human PrimPol obtained from the AlphaFold database (accession code AFQ96LW4) and assuming that Arg417 interacts with a 5’-triphosphate of a primer (Figure 5). The template-primer obtained from the PriL-CTD/DNA-RNA structure (pdb ID 5f0q) was manually fitted into the model using “align to molecule” function in PyMOL, by placing the DNA template into the positively charged groove and the triphosphate close to Arg417. Strikingly, the model revealed high similarity in structural organization of the initiation site in the ZnFn and PriL-CTD, including the histidine and methionine residues that stabilize the template base preceding the initiating base-pair (Figure 5C) and two arginines interacting with a triphosphate (Figure 5D). Thus, despite significant difference in the overall fold and coordinated metals (PriL coordinates the 4Fe-4S cluster), ZnFn and PriL-CTD bind the initiating nucleotide and the template-primer in a similar way. Interestingly, Arg424 cannot interact with a 5’-triphosphate and should not play a structural role (Figure 5D). This fact points to other role of Arg424 in DNA synthesis priming, for example, in CTD interaction with NTD in the initiation complex. Noteworthy, the initiation site of PrimPol is located in close proximity to the zinc-binding site.

**Figure 5.**
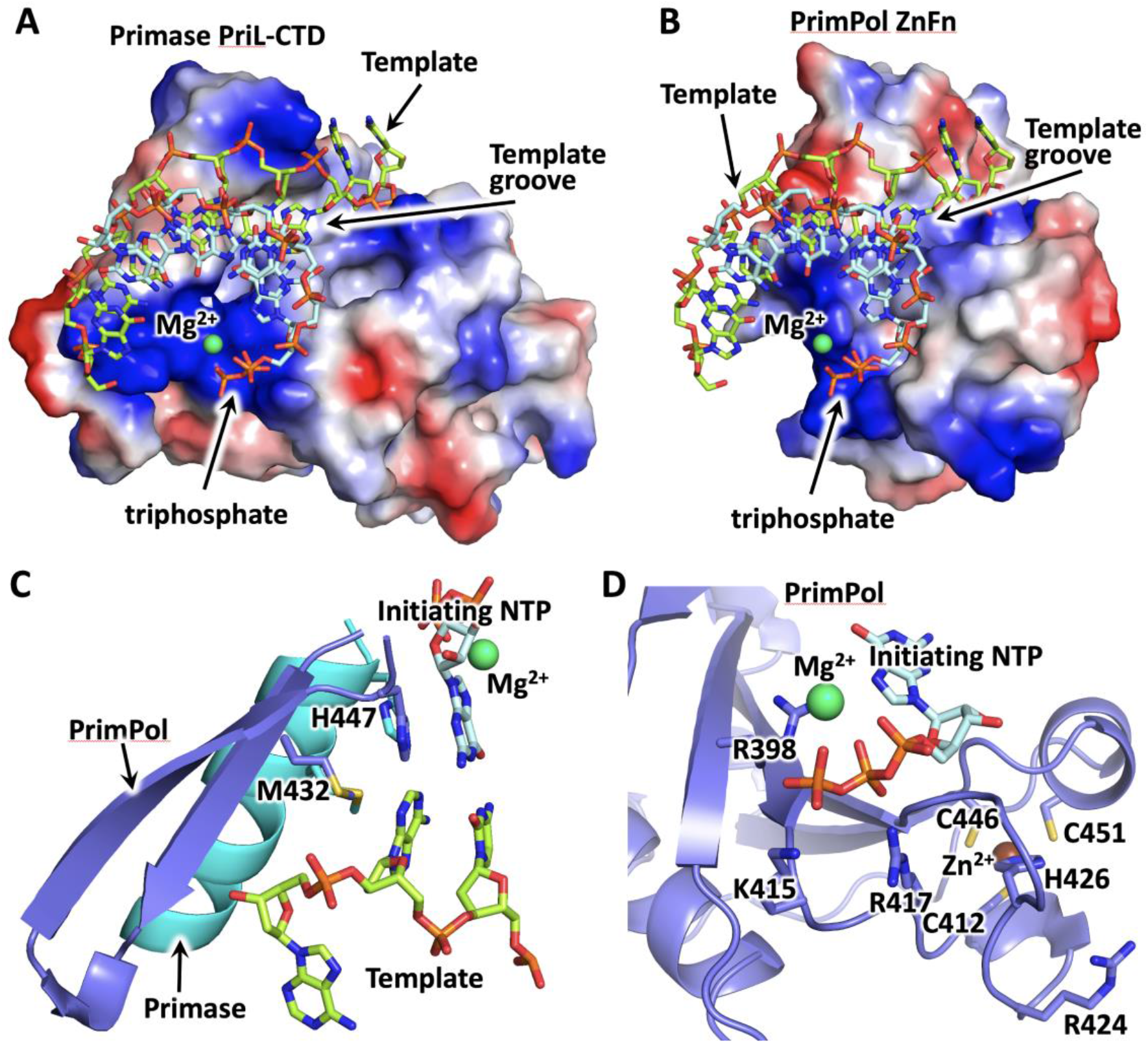
ZnFn and PriL-CTD bind the template-primer and the initiating nucleotide in a similar mode. Comparison of template-primer binding by PriL-CTD (**A**) and ZnFn (**B**) using the coordinates of the PriL-CTD/DNA:RNA complex (PDB ID 5F0Q) and human PrimPol obtained from the AlphaFold database (accession code AFQ96LW4). The protein surface is represented by the vacuum electrostatic potential. **C**. Met432 and His447 of PrimPol stabilize the initiating base-pair and the template base preceding it. The corresponding residues of PriL-CTD have the same position. **D**. The close-up view of the initiation site of PrimPol. DNA, RNA, and amino acids are shown as sticks. PriL-CTD and PrimPol residues colored cyan and slate, respectively. Mg^2+^ and Zn^2+^ ions are shown as spheres and colored green and brown, respectively. The figure was prepared using the PyMOL Molecular Graphics System (version 1.8, Schrödinger, LLC).

### The role of the RBM in primase activity

We demonstrated that the CTD negatively regulates the DNA polymerase activity of full-length PrimPol. Moreover, the RBM of the CTD is involved in modulation of PrimPol binding to DNA with the 5’-triphosphate. PrimPol co-purifies from human cells along with RPA and mtSSB (3). RPA recruits PrimPol to DNA damage sites in cells, and RBM deletion increases cell sensitivity to DNA damaging agents (3). In addition, RPA stimulates the DNA polymerase and DNA primase activities of PrimPol *in vitro* (31, 33). PrimPol binds to the NTD of the RPA1 subunit (RPA70N) (20, 31). On the surface of RPA70N, there is a positively charged region responsible for interaction with many RPA partner proteins (34). The RBM-B motif, on the contrary, contains negatively charged amino acids whose mutations (residues Asn551 and Glu548) disrupt the interaction of PrimPol with RPA. The possible mechanism of PrimPol regulation may include an increase in affinity to DNA by neutralizing the negative charge upon biding with RPA. It is also possible that RPA binding promotes some structural reorganization of the CTD favoring physical interaction between ZnFn and cryptic Gua and/or the triphosphate of initiating ATP. It can be assumed that the DNA polymerase activity of PrimPol on a DNA template with a primer lacking the 5’-triphosphate mainly depends on the DNA-binding properties of the NTD, which is sensitive to CTD inhibition. Therefore, the inhibitory effect of CTD is stronger than its stimulatory effect based on CTD interaction with a 5’-triphosphate.

## Conclusions

Similar to human DNA primase, the results of this study indicate a division of work between the NTD and CTD of human PrimPol: the NTD is responsible for catalysis, and the CTD interacts with the substrates at the initiation and elongation steps of primer synthesis. The *cis*-orientation during *de novo* DNA synthesis, the regulatory role of the CTD in DNA binding, and the requirement of a triphosphate group for stabilization of initiating and elongating complexes of PrimPol with DNA together describe the mechanism of primase activity similar to that seen in human primase.

## DATA AVAILABILITY

The data that support the findings of this study are included in the Supplementary Data file or available from the corresponding author upon request.

### SUPPLEMENTARY DATA

Supplementary Data are available at NAR Online.

## ACKNOWLEDGEMENT

We thank K. Jordan for editing this manuscript.

## FUNDING

This work was supported by the National Institute of General Medical Sciences grant R35GM127085 (THT) and the Russian Science Foundation grant 18-14-00354 (AVM).

### Conflict of interest statement

None declared

## Author contributions

EOB: protein purification, preparation of experiments, data analysis; DIG: preparation of experiments, AGB: design of experiments, data analysis, draft correction; AVM: data analysis, design of experiments, writing the draft of the paper, preparation of figures; funding; THT: data analysis, design of experiments, modeling, preparation of figures, draft correction; funding.

## Notes

### Competing Interest Statement

The authors have declared no competing interest.

