## Supplemental Information for "The role of catalytic and regulatory domains of human PrimPol in DNA binding and synthesis"

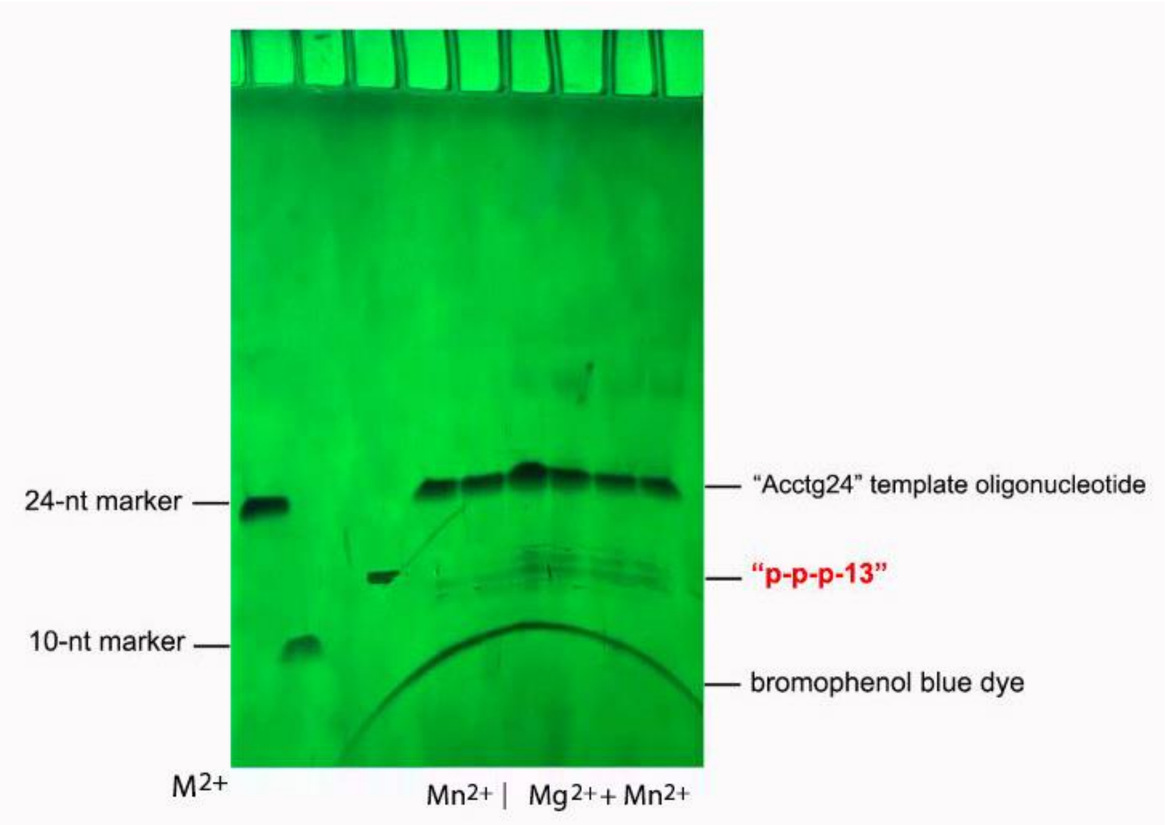


**Figure 1S.** Isolation of the "p-p-p-13" oligonucleotide from the 20% Urea-PAGE. DNA primase reactions with PrimPol and Acctg24 template were carried out in the presence of $Mn^{2+}$ ions or both $Mn^{2+}$ and $Mg^{2+}$ ions. The products of DNA primase reactions were loaded on a gel and visualized by UV shadowing on a fluorescent TLC plate.

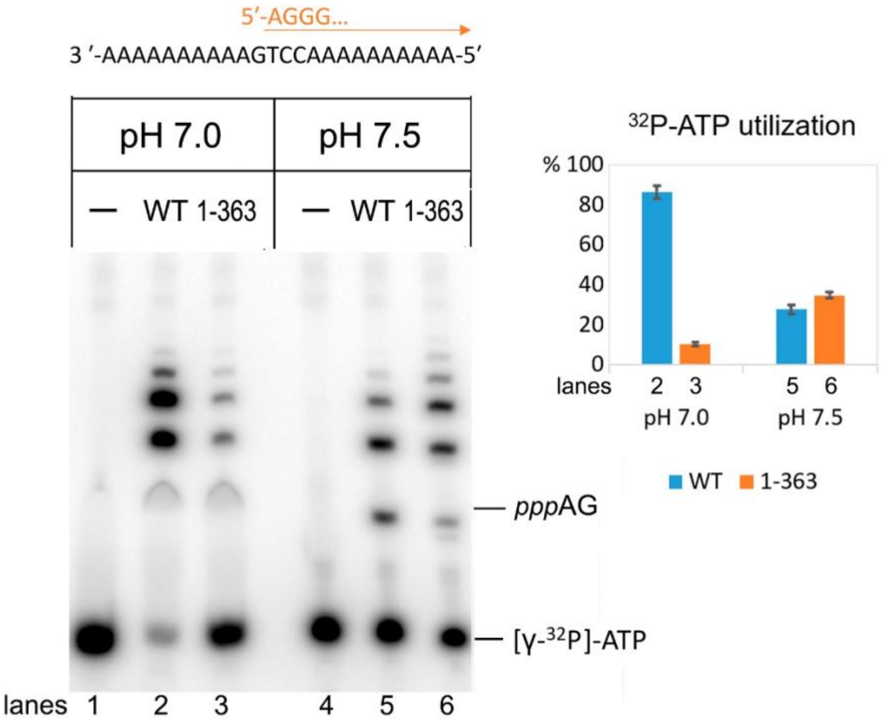


**Figure 2S.** The DNA primase reactions with the full-length PrimPol and PrimPol$_{1\text{-}363}$ variant (NTD) were carried out in the presence of 1 mM $Mn^{2+}$ ions, [γ-$^{32}$P]-ATP, rATP and dGTP at pH 7.0 or 7.5. PrimPol makes an A-G mismatch upon the third and following dGMP insertions.

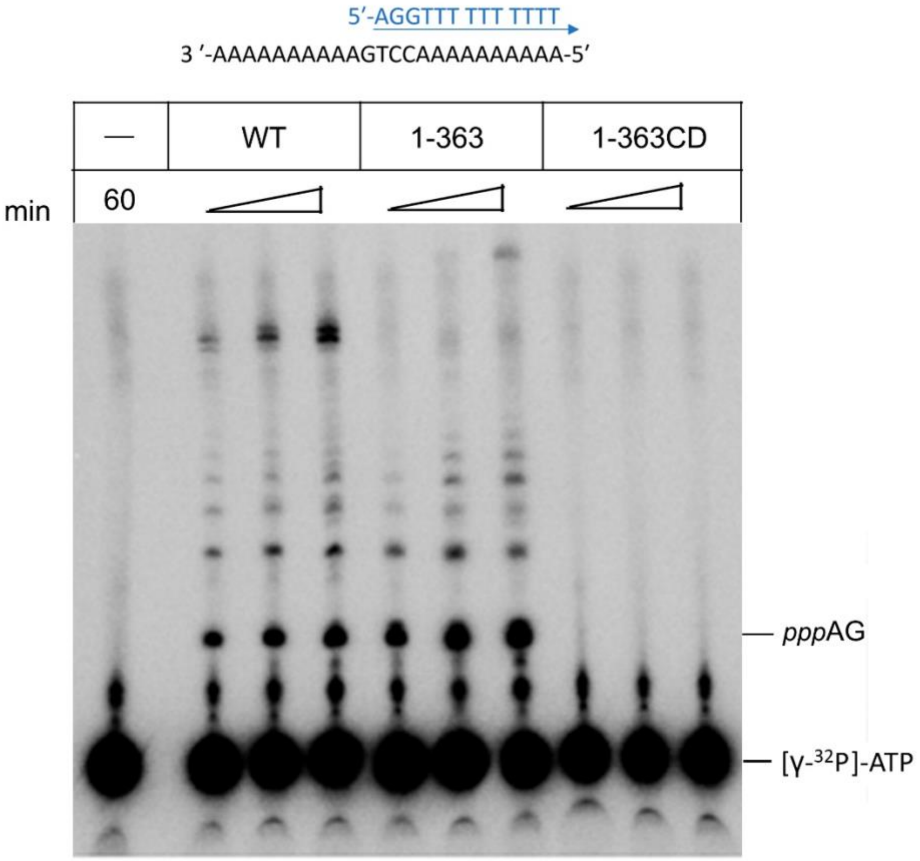


**Figure 3S.** The DNA primase reactions with the full-length PrimPol, $PrimPol_{1-363}$ and $PrimPol_{1-363CD}$ variants were carried out at pH 7.5 in the presence of 1 mM $Mn^{2+}$ ions, [γ-$^{32}$P]-ATP, rATP, dGTP and dTTP; 15 – 60 min incubation time.
